# Identification of novel pulmonary vein nodes as generators of ectopic arrhythmic foci for atrial fibrillation: An immuno-histochemical proof

**DOI:** 10.1101/2021.06.10.447815

**Authors:** T. Gupta, M. Kaur, D. Sahni

**Affiliations:** Department of Anatomy, Postgraduate Institute of Medical Education & Research, Sector-12, Chandigarh-160012, India

**Author notes:** Correspondence to: Dr. Tulika Gupta, Department of Anatomy, Post Graduate Institute of Medical Education and Research, Chandigarh - 160012, India. E-Mail.

**Keywords:** Atrial muscle sleeve, pulmonary vein, specialised cardiac conductive tissue, atrial fibrillation

## Abstract

**Introduction:** The atrial muscle sleeve (AMS) of the pulmonary vein is the most common source of the arrhythmogenic triggers in atrial fibrillation (AF). Anatomical substrate generating these ectopic currents is still elusive. The present study was designed to study the AMS of pulmonary veins with an emphasis on the structural basis which might govern AF initiation and perpetuation.

**Materials and Methods:** The study was conducted on longitudinal tissue section of pulmonary vein, taken from 15 human cadaveric non-diseased hearts. Tissue was studied histologically using H&E and Gömöri trichrome stain. The pacemaker channels were identified by immunohistochemistry using monoclonal HCN4 and HCN1 antibodies.

**Results:** The AMS was identified in each pulmonary vein, located between the tunica adventitia and tunica media. A node like arrangement of myocytes was seen within the AMS in 30% of veins. It had a compact zone limited by a fibrous capsule and contained much smaller, paler and interconnected myocytes. Outside the capsule there was a zone of dispersed, singly placed myocytes separating the compact zone from the working myocytes of the AMS. HCN4 and HCN1 antibodies were expressed on the cell membrane of nodal myocytes, while the working myocytes demonstrated none to minimal staining.

**Conclusion:** Pulmonary veins nodes are similar to the specialized cardiac conductive tissue in, histological arrangement of compact and transitional zones, cellular characteristics, and the presence of pacemaker channels. They might be the anatomical basis of ectopic arrhythmogenic foci. To our knowledge these nodes are being described for the first time in human.

## Introduction

Previous electrophysiological studies have found that “triggers” that start & then maintain atrial fibrillation (AF) originate from the atrial muscle sleeve present in pulmonary vein (PV) wall in 90-95% of cases (Calkins et al., 2018; Gao et al., 2011; Honjo et al., 2003; Wang et al., 2003). The pulmonary veins have demonstrated electrical activity even when they are disconnected from the heart (Jais, 2000; Marrouche et al., 2002; Yamane et al., 2001). Moreover, drug resistant arrhythmia is treated by radiofrequency catheter ablation of pulmonary vein walls or PV antral isolation (Gupta et al., 2020). These facts establish the presence of triggers or specialised cardiac conductive cells in pulmonary vein walls but the anatomical basis of these triggers is not yet understood. The presence of specialized conduction cells in human pulmonary veins in AF patients has been reported (Perez-Lugones et al., 2003). HCN (Hyperpolarisation-activated cyclic nucleotide–gated) channels or pacemaker channels serve as non-selective, voltage-gated cation channels in plasma membranes of the heart. HCN4 plays a key role in the generation and modulation of cardiac rhythmicity (Larsson, 2010). HCN4/PAS-positive cardiomyocytes have been found at the pulmonary vein-left atrium junction in chronic AF (Nguyen et al., 2009). Thus, the presence of automaticity potential in PV is well accepted but its anatomical substrate is still elusive. Therefore, the present study of pulmonary veins was done to understand the structural basis which might govern the initiation & perpetuation of abnormal electrical activity. The entire length of the pulmonary vein was studied histologically in 60 PVs following which an immuno-histochemical study using monoclonal HCN1 and HCN4 antibodies was conducted.

## Material & Methods

### Material

The present cross-sectional study was conducted on 15 human cadavers (48 - 85 years; 10 males and 5 females) procured from the Department of Anatomy, PGIMER, Chandigarh. The study has been conducted on donated cadavers and informed consent of the family was obtained at the time of every donation. The study has been conducted according to the principles outlined in the Declaration of Helsinki. Inclusion criterion: No history of cardiac disease and morphologically normal human hearts. Exclusion criterion: History of cardiac disease &/or cardiac intervention. Hearts with any gross abnormality or pathological changes in valves.

### Method

The thoracic cavity was opened by retracting both halves of the rib cage laterally. The mediastinal contents were extracted out of the cadaver as a single block containing heart, lungs and oesophagus. The heart was separated and both ventricles were removed by incising along the atrio-ventricular groove. The posterior wall of the left atrium was exposed. A longitudinal tissue section was taken along the length of each pulmonary vein, including the posterior wall of the left atrium.

### Histology

Tissue processing and blocking was done. 5μm thick sections of each segment were taken and stained with Haematoxylin & Eosin and Gömöri one step trichrome stain. The Gömöri stain was used to differentiate between myocytes, collagen fibres, and nuclei. Myocytes stain red, collagen fibres appear blue-green, and nuclei stain black.

Gömöri one step trichrome stain: Sections were de-paraffinized and brought to water and were treated with picric acid at 60o for 20 minutes. Excess of picric acid was removed by washing under tap water and then stained with celestian blue for 5 minutes. Distilled water rinsing was followed by haematoxylin staining for 5 minutes. Sections were rinsed in distilled water again and stained with Gömöri stain for 20 minutes. Differentiation was done with 1% acetic acid. Each micro slide was examined to study the histological features of the atrial muscle sleeve in detail.

### Immuno-histochemical study

Tissue sections with 5μm thickness were applied to slides pre-coated with poly-L-lysine. The paraffin sections were deparaffinised in xylene using three changes of 10 minutes each. Sections were hydrated gradually through graded alcohols by washing in 100% ethanol, 70% ethanol, and 50% ethanol for 5 minutes each. Endogenous blocking was done with hydrogen peroxide and de-ionised water (1:9) to quench endogenous peroxidase activity for 20 minutes. The antigen was retrieved by incubating tissue sections for 5-7 minutes in Proteinase K antigen retrieval solution (Abcam, ab64220). Incubation with primary monoclonal antibodies against HCN1 (Novus Biologicals, nbp1-22450, dilution 1μg/ml) and HCN4 (Abcam, ab32675 dilution 1:100) was performed for 2 hours at room temperature followed by an overnight incubation at 4°C. Positive control for HCN1 was normal human cerebellum and normal human heart tissue for HCN4. Negative control was used by omitting the primary antibody. The slides were incubated with HRP-conjugated anti-mouse secondary antibody (Abcam, ab97046, dilution 1:2000) at room temperature for 1 hour. Peroxidase activity was developed in 0.5% 3, 3’-diaminobenzidine till the desired stain intensity developed. Counter staining was done with haematoxylin. Sections were cleared in xylene & mounting medium, i.e., di-n-butyl phthalate in xylene (DPX) was added. The slides were covered with a cover slip and observed under a light microscope.

## Results

The wall of the pulmonary veins is comprised of three distinct layers, an innermost intimal layer of lining epithelium resting on the connective tissue, a middle layer of tunica media and an outer layer of loose adventitia. The atrial muscle sleeve was identified in the pulmonary venous wall by its location - a cardiac muscle bundle running in continuation of the atrial muscle wall located in between the adventitia and media of the venous wall. In all cases it was separated from the tunica media by clearly defined connective tissue.

### Node like Structure in pulmonary veins (PV nodes) (Figs. 1–6)

A node like arrangement of myocytes was seen within the myocardial muscle sleeve in 30% of pulmonary veins. This arrangement can be christened as pulmonary vein nodes (PV nodes). Their presence was observed in all four pulmonary veins with almost equal frequency. The location of the PV nodes was variable in different veins. PV nodes were present from the atrio-venous junction to terminal 1/3^rd^ of the myocardial sleeve. They were never found at the terminal end in any of the cases. They were seen in the adventitial edge of the sleeve and also in between the muscle bundles of atrial muscle sleeve but never towards the tunica media. In one case two nodes like structures were found close to each other embedded in the adventitia near the atrio-venous junction (Figure 4).

PV nodes were round to oval in shape and variable in size. The nodes were well circumscribed and limited by fibrous sheet, and were always associated with one or two vessels lying just outside this fibrous capsule. Each node was separated from the normal working myocytes by a distinct transitional zone. Thus, PV nodes can be divided into compact zone, transitional zone, and working myocytes, present from inside to outside of the node (Figs. 1, 2). **Compact zone:** Multiple myocytes were seen within the PV nodes. These myocytes were much smaller, about 1/3^rd^ to 1/5^th^ the size of the working myocytes. The myocytes were longitudinal in shape and at times appeared to have tapering ends. They had a large central oval nucleus with prominent nucleoli. The cytoplasm was scanty and very pale in comparison to the working myocytes. Another striking difference was paucity of striations compared to the working myocytes, although a few red striations could be visualised in Gömöri stained slides (Figure 4). The myocytes were interconnected forming a network like arrangement. Variable amount of connective tissue was interspersed with the myocytes. Multiple small size blood vessels were seen within the nodes. **Transitional zone:** Discrete muscle fibres were seen outside the fibrous capsule. They were circumferentially arranged around the node as a mantle which was 1-4 cells thick. These fibres were singly present, not connected to each other. The fibres were slightly bigger than the nodal myocytes but were smaller than the working myocytes and demonstrated distinct tapering ends. They were almost as pale as the nodal myocytes with no striations and with distinct central nucleus. In most of the cases, this mantle was separated from the capsule by some space.

**Figure 1:**
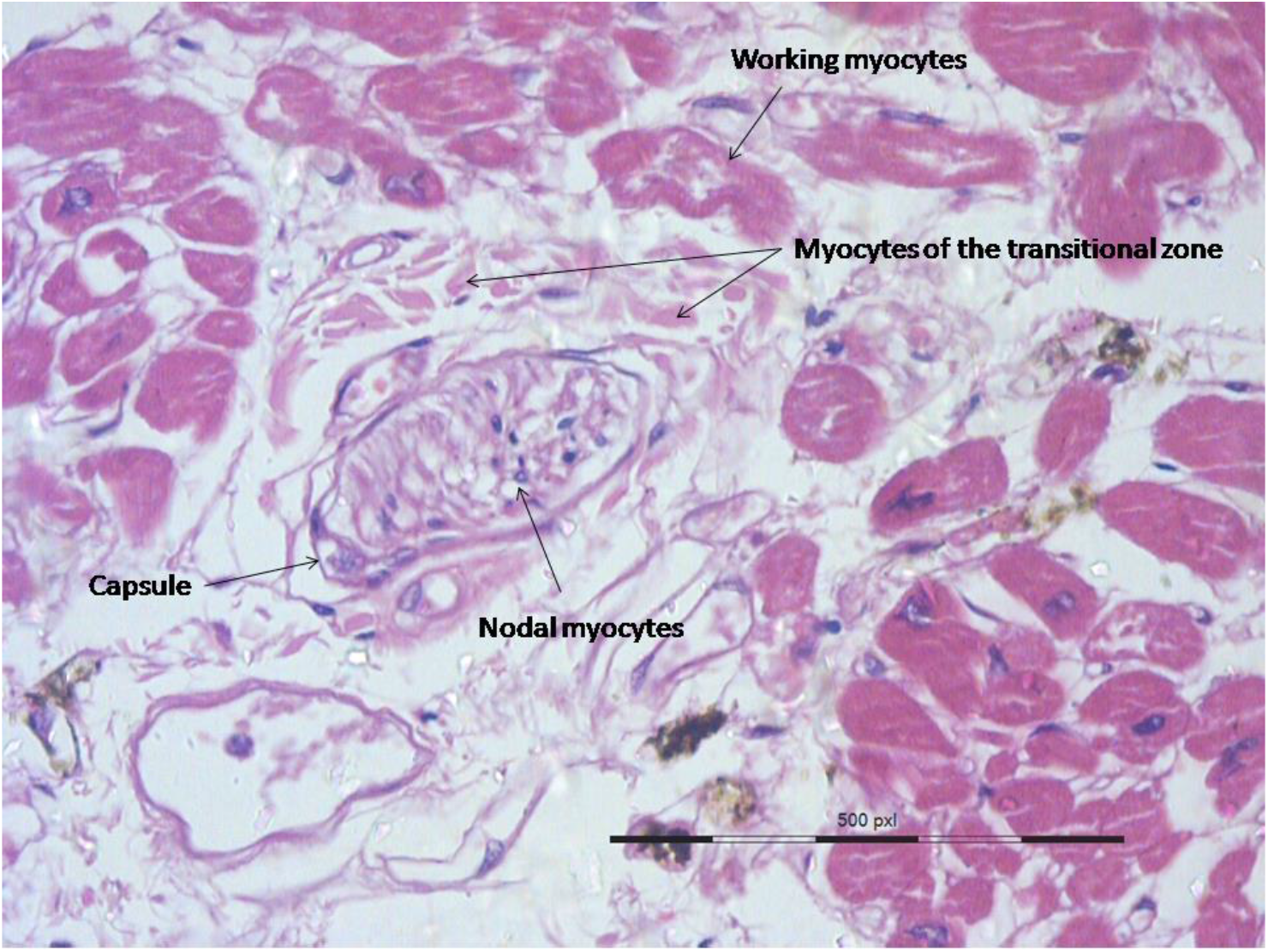
A node like structure is seen lying within the cardiac muscle bundles of the atrial muscle sleeve. Compact part of the node has smaller and paler myocytes with no discernible myofibril forming a syntitium. They are limited by a well-defined capsule. Transitional zone is seen outside the capsule. Transitional myocytes are smaller and much paler than the surrounding working myocytes of the atrial muscle sleeve. Many capillaries are seen around the node. H&E stain at 40x.

**Figure 2:**
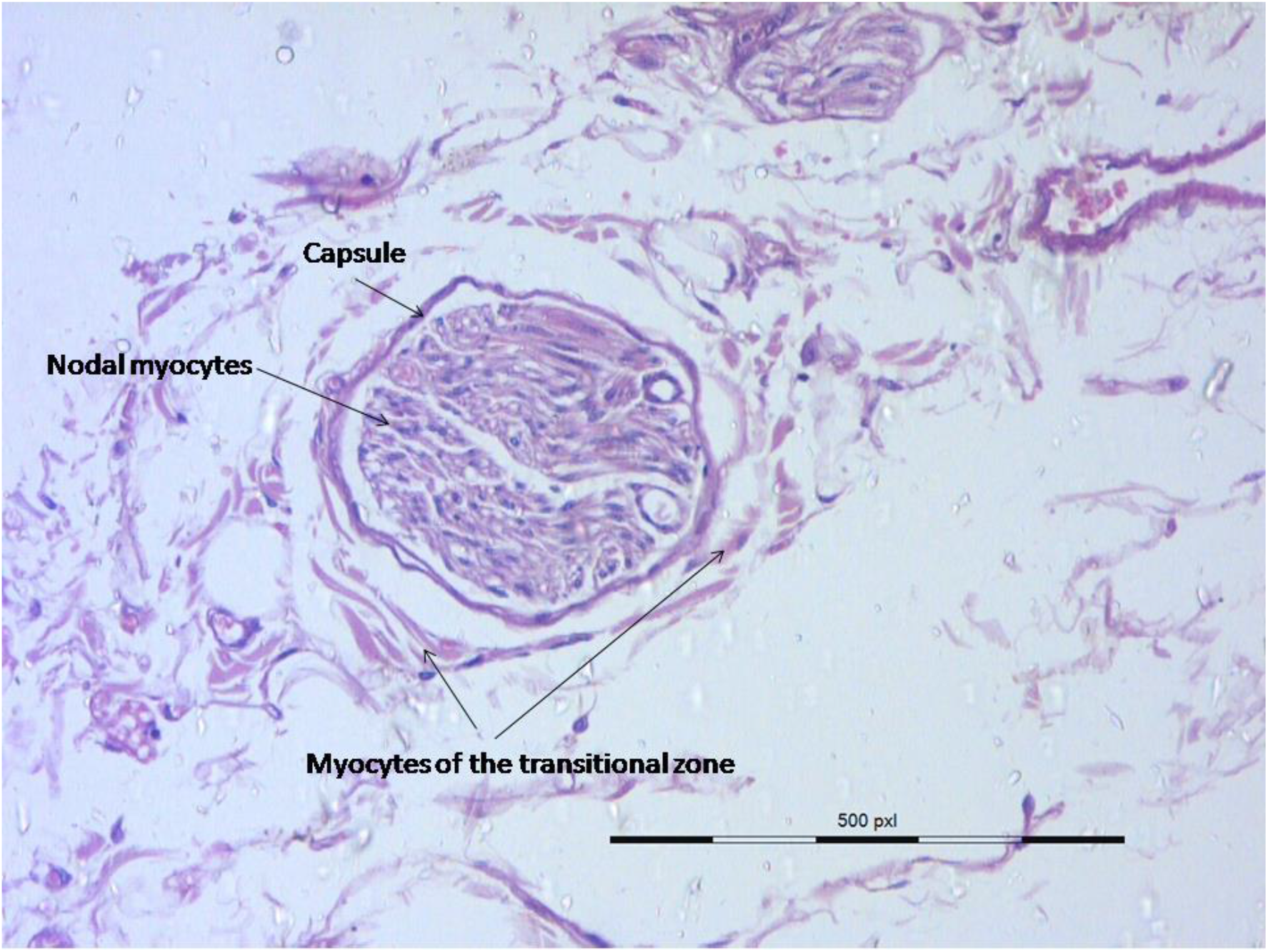
A smaller PV node present in the tunica adventitia near the atrio-venous junction. Capillaries are seen within the capsule. H&E stain at 40x. PV – pulmonary vein

### Working myocytes

These were the myocytes of the atrial muscle sleeve and were arranged in the form of syncytium. In comparison to the myocytes of the compact and transitional zones, the working myocytes were darkly stained. The myocytes were much larger with a central nucleus and distinct striations could be seen.

### Immunohistochemistry(Figs. 5,6)

The myocytes present within the PV nodes demonstrated distinct staining with the monoclonal HCN4 and HCN1 antibodies. The HCN4 and HCN1 immunoreactivity was present at the cell membrane of these myocytes. Very mild presence was also detected in the cytoplasm in few myocytes. The myocytes of the transitional zone also demonstrated HCN immunostaining. On the other hand, the surrounding working myocytes demonstrated none to minimal staining and none for HCN4 and HCN1. HCN4 immunoreactivity when present in the working myocytes was more in the sarcomeres present in the cytoplasm than on the cell membrane.

## Discussion

There is a substantial body of evidence which establishes that focal points of enhanced automaticity are present on pulmonary veins. Pulmonary veins are considered to be the most common source of arrhythmogenic focus in atrial fibrillation. These have been proven by many electrophysiological studies (Jais, 2000; Marrouche et al., 2002; Yamane et al., 2001). There is substantial clinical data proving that pulmonary veins have the capacity to generate aberrant electric currents; 90-95% of atrial fibrillations are triggered by foci located on the atrial muscle sleeve present in the walls of pulmonary veins. Radiofrequency catheter ablation of these triggers points is the preferred treatment for atrial fibrillation with drug resistance and is considered the first line of treatment in paroxysmal atrial fibrillation (Gupta et al., 2020). Despite this overwhelming evidence, the anatomical basis of this focal automaticity remains elusive. In the present study, we have identified circumscribed areas in the PV atrial muscle sleeve which structurally fulfil the criterion for specialised conductive tissue on the basis of their morphological structure examined by histology. Moreover we have demonstrated presence of HCN4 and HCN1, the major constitutive subunit of f-channels in pacemaker cells, on these myocytes (DiFrancesco, 2010; Mitrofanova et al., 2014). We have named these areas PV nodes. PV nodes might be the anatomical substrate of the enhanced automaticity seen in the pulmonary veins. To best of our knowledge these nodes are being described first time in human pulmonary veins.

Criteria for histological diagnosis of specialised conduction tissue include that cardiomyocytes should be histologically discrete, should be traceable within serial histological sections and most importantly, should be insulated from the adjacent myocardium (Anderson et al., 2016; Aschoff, 1910; Monckeberg, 1910). The PV nodes described by us fulfil all three criteria. These nodes were discrete, circumscribed and clearly demarcated from the surrounding tissue by a well-defined fibrous capsule (Figs. 1–6). The histological features of these nodes closely resemble those of the S-A node and the A-V node. The S-A node and the A-V node have cells of varying morphology which are small, pale staining and have none to low myofibrils (Mitrofanova et al., 2014; Randhawa et al., 2017; Vassall-Adams, 1983). Waller (1993), has described the nodal cells as meshwork of pale cells, which are interwoven with collagen and elastic fibres (Waller et al., 1993). The cells present in nodes of the pulmonary veins in the present study were smaller and much paler then the surrounding myocytes of the atrial sleeve (Figure 1). Nodal cells were of variable morphology from cylindrical, spindle like to circular shaped, and displayed a large central nucleus (Figs. 4, 5a). These myocytes had minimal striations in comparison to working myocytes (Figure 1). These cells also displayed interconnections forming a network-like arrangement (1, 2, 6). The PV nodes also contained connective tissue (Figure 4) interspersed with myocytes. Hence, these PV nodes were populated by the cells which demonstrated features very similar to that of specialised node cells in the S.A. node. All the PV nodes were surrounded by a well-defined fibrous capsule. Similar to the transitional zone of the A-V node (Randhawa et al., 2017), these PV nodes also exhibited a mantle zone of myocytes. The mantle myocytes were bigger than the nodal myocytes but smaller than the working myocytes, separating the nodal from the working myocytes (Figs. 1, 2). Thus, they resemble the myocytes found in the transitional zone of the A-V node (Randhawa et al., 2017; Ross and Pawlina, 2011; Waller et al., 1993). Moreover, these myocytes were present singly and a bit scattered, quite similar to the description given by Rossi (1980) that the transitional zone is not a definite layer external to the compact atrio-ventricular node but is made up of thin separate fascicles (Rossi, 1980).

**Figure 3:**
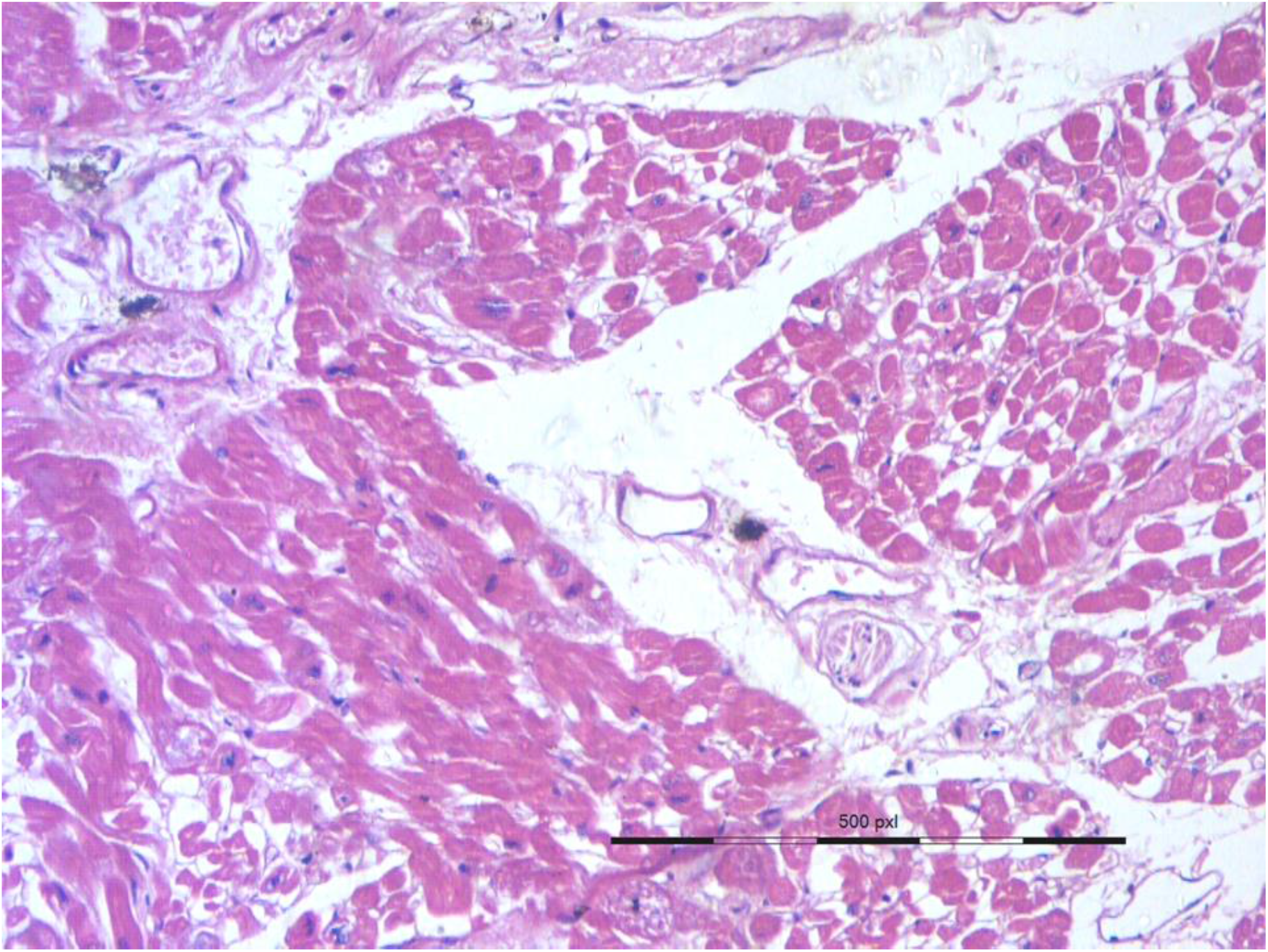
A small PV node lying within the cardiac muscle bundles of the atrial muscle sleeve. H&E stain at 40x. PV – pulmonary vein

**Figure 4:**
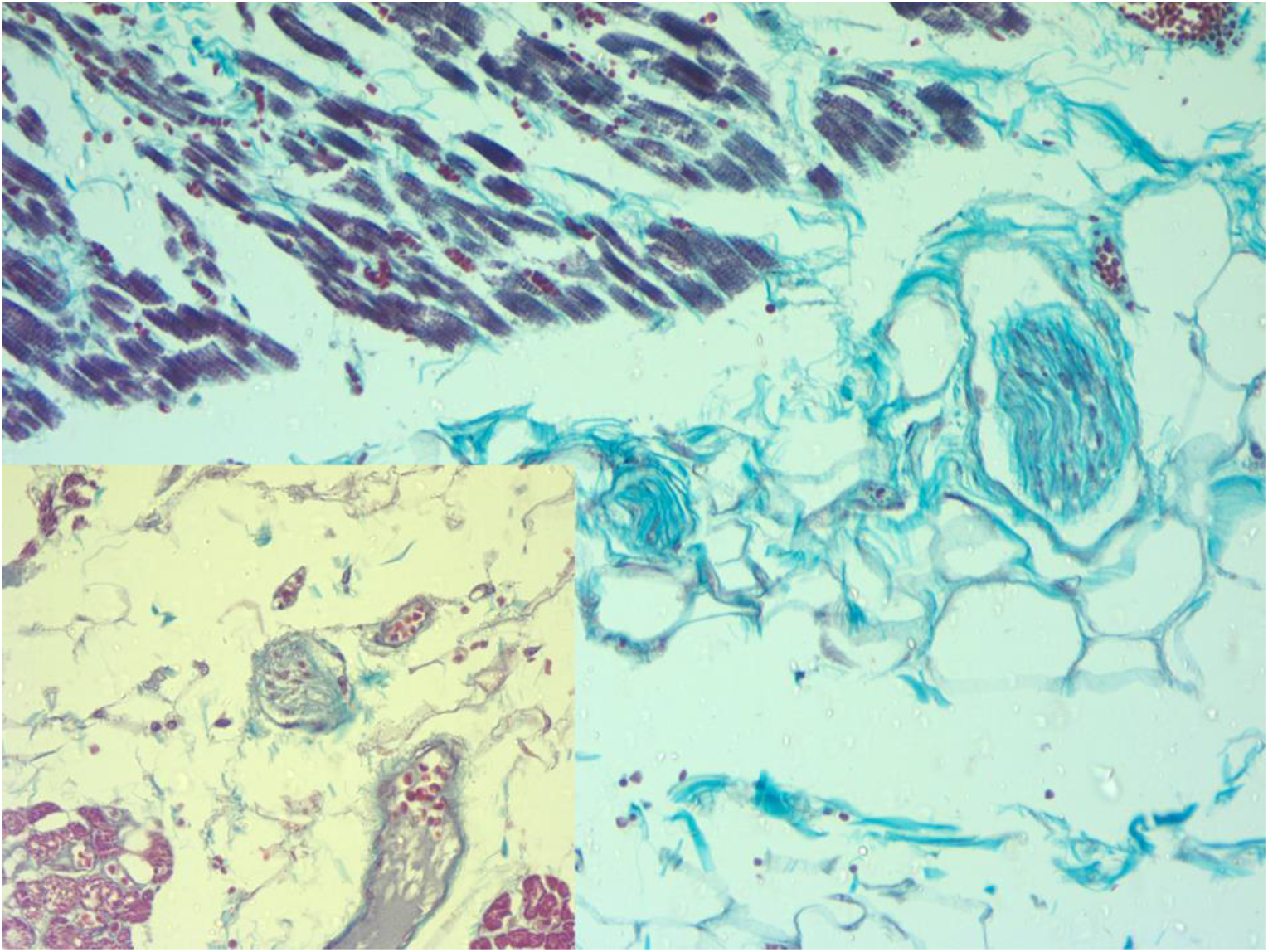
Gömöri one step trichrome stain 40x. Two nodes are seen lying in adventitia. The working myocytes of the atrial sleeve are seen in the upper field. They have taken a red colour because of an abundance of myofibrils in the cytoplasm. Myocytes of compact zone have no myofibrils in the smaller node but a few myofibrils (red) can be seen in the bigger node. Inset shows another case. A PV node with small myocytes is seen in the centre while working myocytes are seen in three corners. PV – pulmonary vein

**Figure 5:**
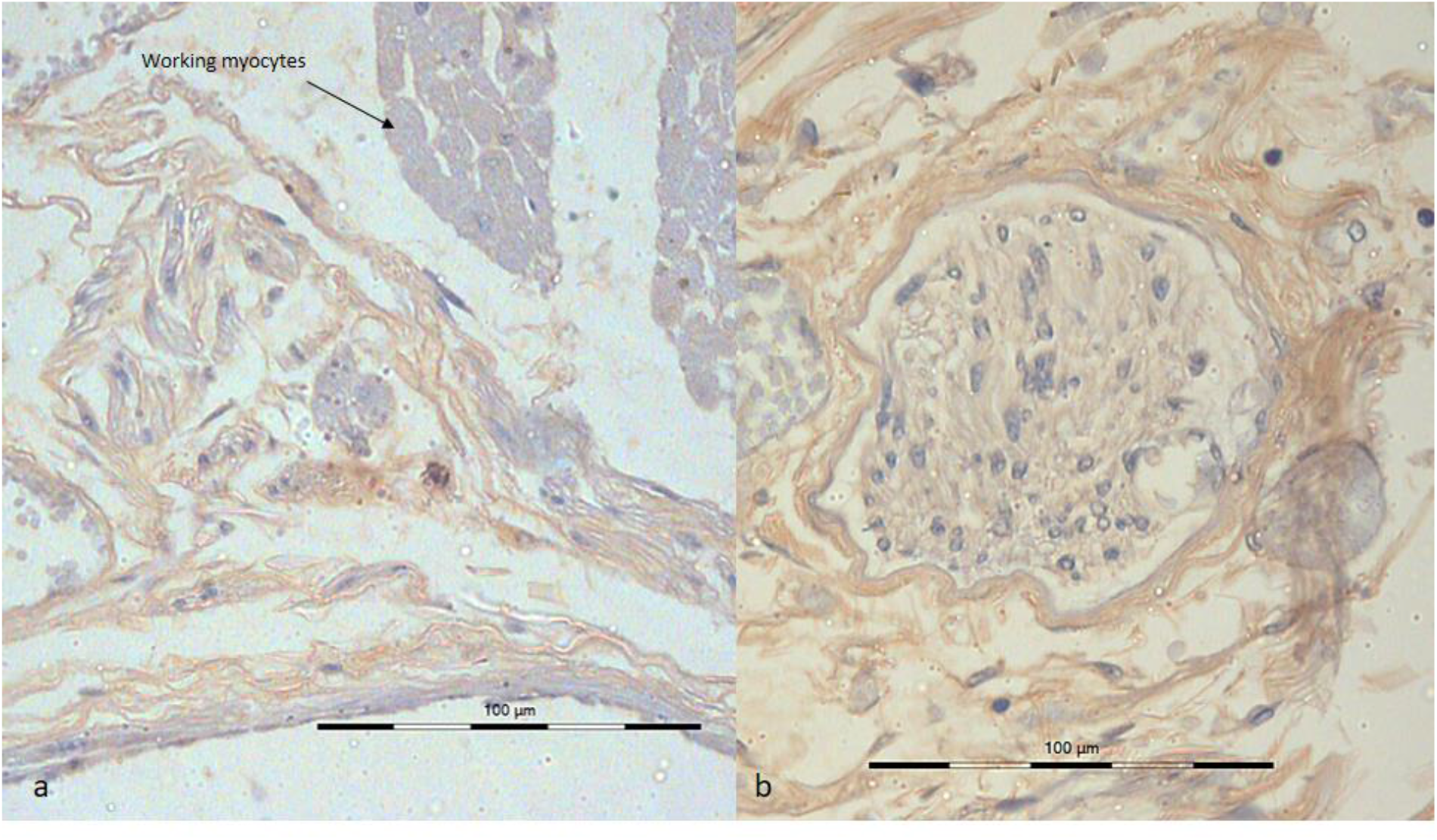
Immunohisto-chemistry using monoclonal HCN4 antibody at 40x. HCN4 immunoreactivity on cell membrane is seen in compact as well as transitional myocytes of the PV node. In contrast the working myocytes seen in the figure a. show none to very mild staining with HCN4 antibody. HCN (Hyperpolarisation-activated cyclic nucleotide–gated) channels, PV – pulmonary vein.

**Figure 6:**
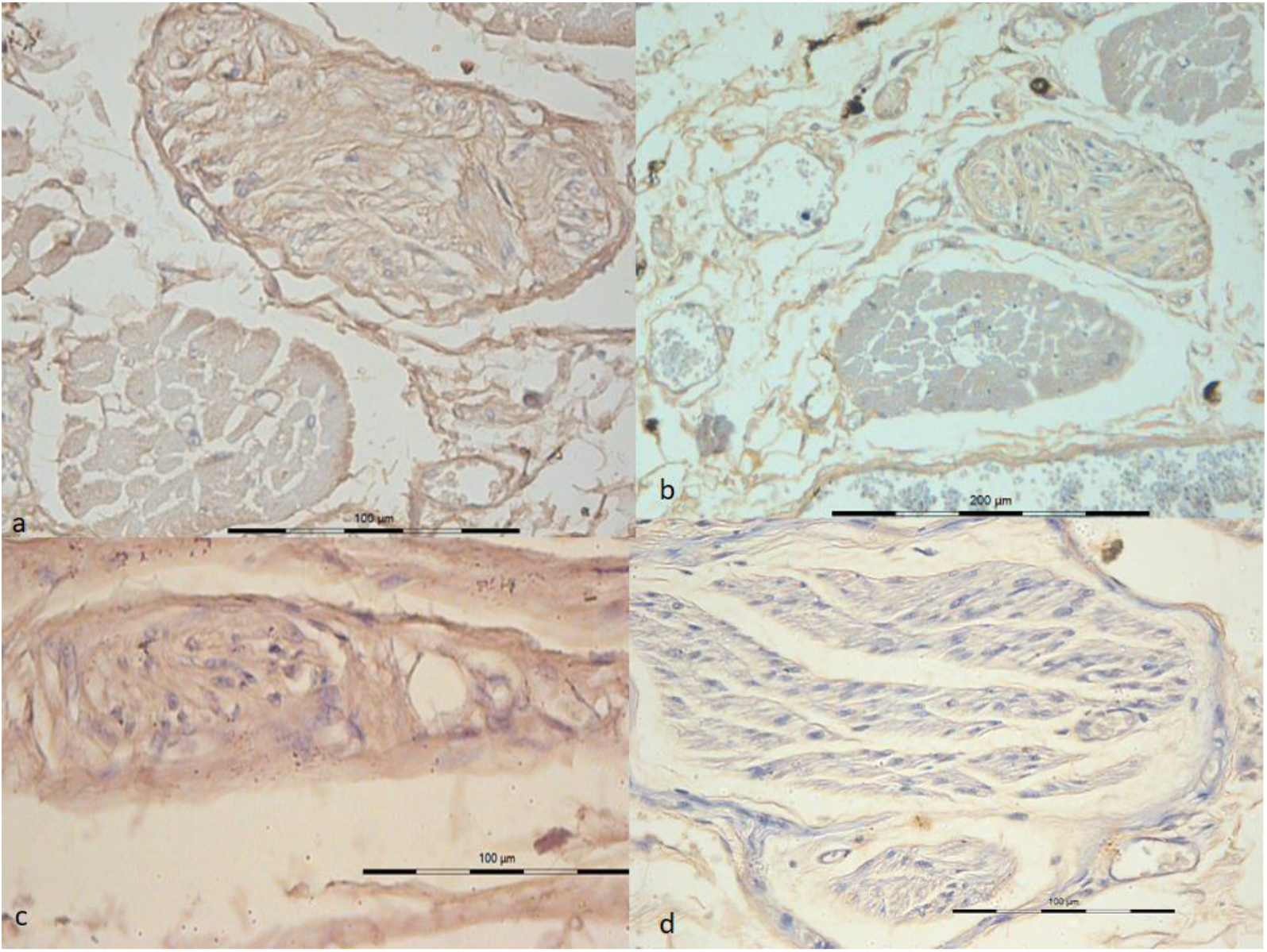
Immunohisto-chemistry using monoclonal HCN4 and HCN1 antibodies. a. and c. show immunostaining by HCN1 antibody at 40x, while b. (20x) and d. (40x) show immunostaining by HCN4 antibody on PV nodes. HCN (Hyperpolarisation-activated cyclic nucleotide–gated) channels.

Previous studies have attempted to identify these specialised myocardial cells responsible for ectopic foci on pulmonary vein. Their reports are at variance. Masani (1986) described node like cells present either singly or in clusters in the atrial muscle sleeve in rats; these were smaller than ordinary myocytes and their cytoplasm lacked myofibrils (Masani, 1986). Thus, these clusters of node-like cells are similar to the nodal cells of the present study. It was observed that these node-like cells may have the potential for pace making activity and represent an ectopic pacemaker centre in the pulmonary vein. Perez-lugones et al. (2003) reported the presence of specialized conduction cells in human pulmonary veins in AF patients. They demonstrated P cells, transitional cells, and purkinje cells in the PV myocardial sleeves. But they did not find the specialised conduction cells in control patients or might have missed them as of their own admission (Perez-Lugones et al., 2003). Nguyen et al. (2009) described a complex histopathological substrate characterized by HCN4-/PAS-positive cardiomyocytes and interstitial Cajal-like cells scattered in the fibrotic tissue, inflammatory infiltrate, and sympathetic nerve structures in the atria and the PV muscle sleeves of chronic AF patients (Nguyen et al., 2009). On the other hand, various authors (Kholová and Kautzner, 2003; Saito et al., 2000; Tagawa et al., 2001) have not found such specialised conductive cells in the pulmonary veins. However, the authors have acknowledged that adequate differentiation of these cells would require detailed electron microscopy and/or histochemical study (Kholová and Kautzner, 2003).

We identified these specialised myocardial cells by light microscopy using special stains and further strengthened the results by immuno-histochemistry. We have identified presence of pace maker channels HCN4 and HCN1 on the PV node myocytes using monoclonal antibodies. Hyperpolarization-activated cyclic nucleotide–gated (HCN) channels or pacemaker channels are integral membrane proteins that serve as non-selective voltage-gated cation channels in the plasma membranes of heart. HCN4 plays a key role in the generation and modulation of cardiac rhythmicity as they are responsible for the spontaneous depolarization in pacemaker action potentials in the heart (Larsson, 2010; Nguyen et al., 2009). HCN channels are present mainly in cardiac conduction system; they are highly expressed in the sinus node (SAN) (Li et al., 2014). Li et al. (2015) detected that all three cardiac HCN isoforms (HCN1, HCN2 and HCN4) had higher expression in the SAN than the atria (Li et al., 2015). HCN1 was almost exclusively expressed in the SAN. The HCN4 channels are responsible for the hyperpolarization activated funny current (If) essential to SAN automaticity. Mutations in the HCN4 gene are known to cause rhythm disturbances, such as inherited sinus bradycardia or familial sinus node disease (DiFrancesco, 2010; Lubitz et al., 2009). Thus, detection of HCN1 and HCN4 based channels on the PV nodes in the present study (Figs. 5, 6) further substantiates the claim that these nodes are populated by specialised cardiac conductive tissue which has the potential to generate pacemaker currents. Increased expression of HCN4 on the pulmonary veins of aged dogs has also been reported (Li et al., 2012).

The embryological basis of the presence of specialised conductive tissue in the pulmonary veins can be explained. In developing hearts, the atrio-ventricular node appears in the dorsal endocardial cushion of the atrio-ventricular canal at about the 6th week of intra-uterine life. Thus, endocardial cushions, which are derived from the neural crest cells, are involved in the development of the specialised cardiac conduction tissue. Developmentally, the posterior smooth part of the left atrium is formed by absorption or incorporation of endocardial cushions of four pulmonary veins (Schoenwolf et al., 2015). The posterior wall of the left atrium shares neural crest origin with the A-V node and it can be hypothesised that they have the potential capacity to form specialised conductive tissue. The atrial muscle sleeve of the pulmonary veins is a layer of cardiac muscle which extends from the posterior wall of the left atrium onto the pulmonary veins (Gupta et al., 2020). The PV nodes in the present study have been detected within this atrial muscle sleeve. A murine model was used to delineate the developing cardiac conduction system by Jongbloed et al. (2004). They found the cells positive for CCS-lacZ, indicative of the developing conduction system, in pulmonary veins. They postulated that these might become active to be arrhythmogenic in a diseased state (Jongbloed et al., 2004).

The PV nodes found in the present study are morphological similar to the specialised cardiac conductive tissue at the cellular level as well as in the histological arrangement. The nodal myocytes demonstrate the presence of pacemaker channels. Thus, these PV nodes have the structural capability for pace making activity. These PV nodes might be the focal triggers of enhanced automaticity in the pulmonary myocardial sleeve. This trigger is probably propagated because of the morphological features of the cardiac muscle sleeves. In one of our previous studies, we reported the presence of long, oblique and spirally arranged fibre bundles denoting that lots of crossing over or twisting of the cardiac fibres is present in the muscle sleeve. Many areas denoting angular change in the direction of the fibres were also noted (Gupta et al., 2020). All these morphological features predispose the muscle fibres to propagate and promote the aberrant electrical current by providing significant substrates for re-entry (Tan et al., 2006). We also found typical terminal postsynaptic autonomic ganglion in the atrial muscle sleeve of the pulmonary veins (Gupta et al., 2020) which belong to the intrinsic autonomic nervous system (ANS) of the heart. Hyperactive ANS may have important contributions in the focal AF of PV origin (Calkins et al., 2018).

Thus, the anatomical substrate capable of generating enhanced automaticity is present in the normal heart. The conversion of these potential pacemakers, into arrhythmogenic foci might occur due to a precipitating factor like inflammation, oxidative stress etc. which are considered risk factors for AF (Kirklin/Barratt, 1993; Saito et al., 2000). How a potential focus is activated is the missing link in our current understanding. Whether PV nodes are present since birth or develop secondarily also needs further investigation.

### Limitations

We have not done the electrophysiological and ultra structural evaluation of the PV node. A similar study on pulmonary veins obtained from known AF patients is also required.

